# Fungal VOCs regulate the expression of WOX5 at the root meristem of plants under osmotic stress in Arabidopsis and *Populus tremuloides* seedlings

**DOI:** 10.1101/2025.09.17.676813

**Authors:** Esperanza Miñambres, Jorge Señorans, Oscar Lorenzo, Mónica Calvo-Polanco

**Affiliations:** Institute for Agribiotecnology Research (CIALE), Department of Botany and Plant Physiology, University of Salamanca, C/del Duero, 12, 37185 Villamayor, Salamanca, Spain

**Keywords:** volatile organic compounds (VOCs), WOX5, abiotic stress, poplar, arabidopsis

## Abstract

Communication between plants and fungi occurs even before direct physical contact via various chemical signaling pathways, among them, volatile organic compounds (VOCs). Fungal VOCs, including those emitted by ectomycorrhizal and symbiotic endophytic fungi, significantly influence plant development and root architecture. However, the molecular signaling mechanisms underlying its regulation remain unknown. The quiescent center (QC), located within the primary root meristem, serves as a reservoir of stem cells that generate daughter cells and constituting a key regulatory site for architectural modulation. These daughter cells either divide or differentiate to form the transition zone, maintaining a crucial balance for root meristem homeostasis. WOX5, a homeobox transcription factor expressed predominantly in the QC, regulates surrounding stem cells by repressing their differentiation, thereby preserving the stem cell niche and is essential for root meristem integrity. In this study, we aimed to analyze the effects of fungal VOCs emitted by *Laccaria bicolor, Hebeloma cylindrosporum*, and *Serendipita indica* on the root architecture and meristem regulation in *Arabidopsis thaliana* and *Populus tremuloides*. Our results demonstrate the impact of these fungal VOCs on the root architecture of both species and underline the importance of WOX5 regulation mediated by fungal VOCs for maintaining root meristem homeostasis.

## Introduction

Volatile organic compounds (VOCs) are molecules emitted by plants, microbial and fungal communities. They are responsible for the promotion of main changes in plant development, including root architecture conformation and lateral root formation. (El Jaddaoui et al., 2023; Inamdar et al., 2020). Our previous study on the role of the VOCs emitted by *Laccaria bicolor, Hebeloma cylindrosporum* and *Serendipita indica* in Arabidopsis plants (Miñambres et al., unpublised), demonstrated the role of the VOCs of those fungi in the enhancement of plant development and lateral root formation through their action in the auxin/cytokinin pathway both under control and osmotic stress treatments. Other studies have highlighted the positive impact of fungal VOCs on plants, including promotion of growth and root development, and enhanced photosynthesis capacity (Venneman et al., 2020, Ditengou et al., 2015). These beneficial effects are particularly determinant in the context of plant defense against various abiotic (Fraj & Werbrouck, 2023; Li & Kang, 2018; Jalali et al., 2017) and biotic stresses (Kumar et al., 2023; Kumar et al., 2021; Singh et al., 2021; Li & Kang, 2018), although the main mechanisms controlling those processes remain elusive. Fungal volatile organic compounds have been also shown to significantly influence root architecture in plants. Specifically, VOCs emitted by *Laccaria bicolor* enhance lateral root formation in *Arabidopsis thaliana* and *Populus tremula × alba* by modulating auxin transport pathways (Ditengou et al. 2015; Felten et al., 2010). Similarly, VOCs and CO_2_ produced by *Serendipita* spp. promote root system development, increasing primary and lateral root growth and density. These effects are mediated through complex hormonal signaling networks involving auxin and cytokinins (Venneman et al., 2020). Collectively, these findings highlight the critical role of fungal VOCs in shaping root architecture, so understanding these volatile-mediated interactions is essential for leveraging fungal VOCs in sustainable agriculture and forestry crops.

Ectomycorrhizal fungi establish symbiotic relationships primarily with woody plants from families such as Pinaceae, Fagaceae, Betulaceae, and Myrtaceae, encompassing around 20,000 species mostly within Basidiomycota and some Ascomycota (Cho et al., 2021; Drijber & McPherson, 2021). Among them, *Laccaria bicolor* associates with temperate and boreal forest trees, including conifers (*Pinus strobus, Pinus pinaster*) and broadleaf species (*Populus trichocarpa, Populus tremuloides*) (Cho et al., 2021; Cope et al., 2019). *Hebeloma cylindrosporum* is commonly associated with pine and holm oak forests across Europe and the Mediterranean (Marmeisse et al., 2004). Due to their ease of culture and genome availability, both fungi serve as model organisms for studying ectomycorrhizal symbioses with trees (Ruytinx et al., 2021; Martin et al., 2008; Marmeisse et al., 2004). On the other hand, endophytic fungi colonize various tissues of terrestrial and some aquatic plants, promoting growth and enhancing resistance to diverse abiotic and biotic stresses (Gautam & Avasthi, 2019). Exhibiting extensive biological diversity, these fungi have been isolated from nearly all vascular plant groups across environments ranging from deserts and tundra to mangroves and forests predominantly from the phylum Ascomycota (Gautam & Avasthi, 2019). *Serendipita indica* is a well-studied endophytic symbiont known for its growth-promoting properties and ability to enhance nutrient uptake and stress tolerance in both monocots and dicots. Its broad host range and ease of laboratory cultivation make it a valuable model for plant-fungus interaction studies (Saleem et al., 2022; Varma et al., 1999).

The root meristem can be divided into the primary meristem (PM), the transition zone (TZ), and the elongation and differentiation zone (EDZ). The quiescent center (QC), located within the primary meristem, gives rise to stem cells that generate daughter cells, which subsequently either divide or differentiate to become part of the transition zone. Therefore, the balance between division and differentiation rates is essential for the proper maintenance of the root meristem (Kong et al., 2018; Stahl & Simon, 2005; Su et al., 2011). WUSCHEL RELATED HOMEOBOX 5 (WOX5) is a homeobox transcription factor predominantly expressed in the QC, where it regulates surrounding stem cells by repressing their differentiation. Inhibition of WOX5 induces differentiation of columella stem cells or distal cells into starch-accumulating columella cells, whereas increased WOX5 expression leads to the over-proliferation of these distal stem cells. Thus, regulation of distal stem cell differentiation is controlled by WOX5 abundance to prevent premature differentiation. WOX5 also suppresses divisions within the quiescent center, maintaining its quiescence by repressing cyclin activity, thereby preserving the QC as a stem cell reservoir essential for root meristem integrity (Burkart et al., 2022; Pardal & Heidstra, 2021; Sarkar et al., 2007).

The objective of this study was to determine the effect of fungal VOCs in the expression of WOX5 under control and osmotic stress conditions and if this effect is not only determinant in Arabidopsis roots development but in *Populus tremuloides*.

## Results

### Volatile organic compounds (VOCs) emitted by *L. bicolor, H. cylindrosporum*, and *S. indica* exert a positive effect on the root development of Arabidopsis

Plants exposed to volatile organic compounds (VOCs) emitted by the three fungal species used in this study exhibited similar effects on the Col-0 *Arabidopsis* seedlings. Phenotypically, seedlings displayed enhanced growth in the presence of fungal VOCs compared to their absence (Figure 1A). Overall, seedlings were larger, as reflected by an increase in fresh weight (Figure 1B), with the effect being most pronounced in response to VOCs from *S. indica*. Furthermore, root analysis revealed an increase in primary root length (Figure 1C) and a higher number of lateral roots (Figure 1D) when exposed to VOCs from any of the three fungi, relative to controls without VOC exposure.

**Figure 1.**
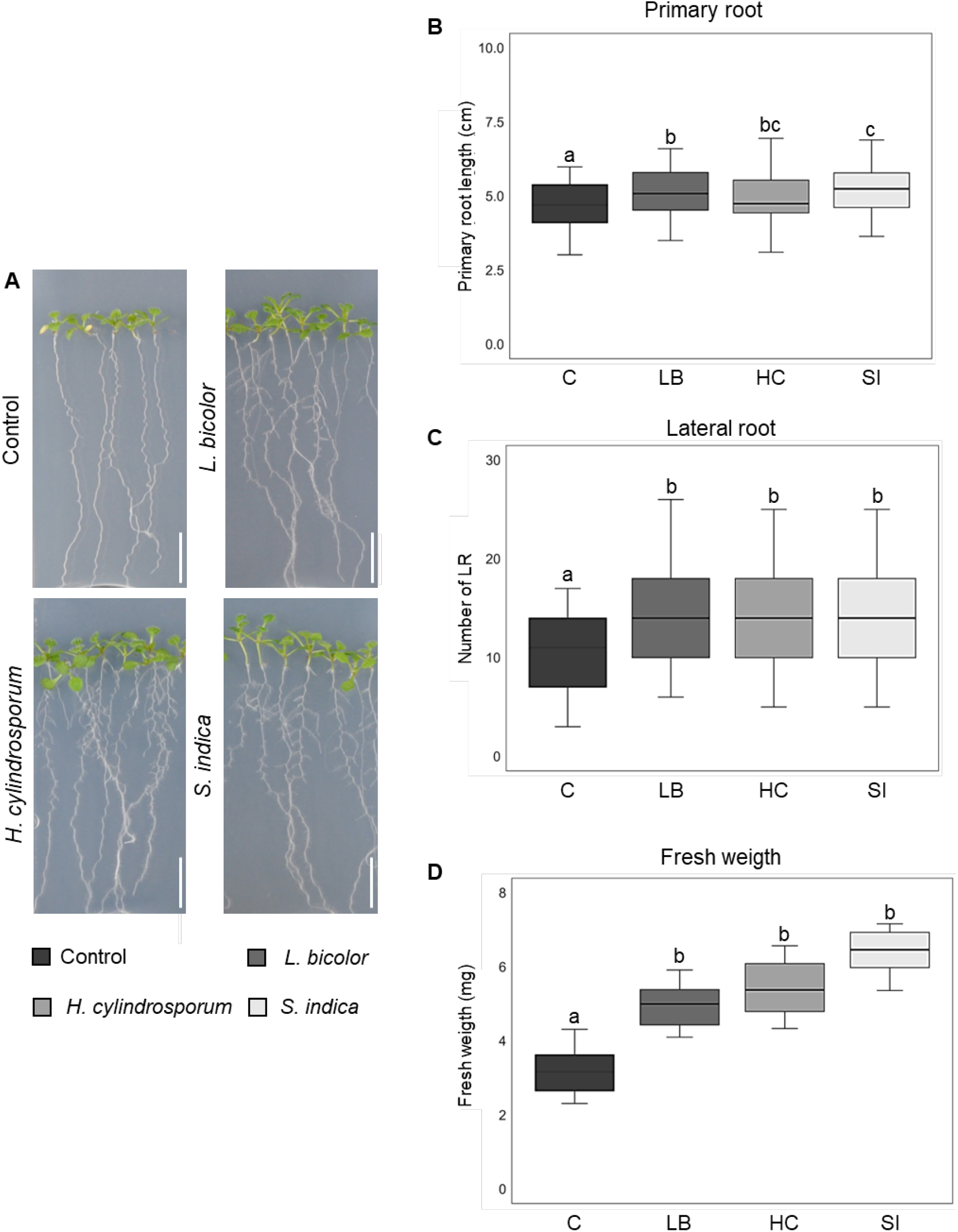
Analysis of the development of 10-day-old Col-0 seedlings in the absence (C) and presence of VOCs emitted by *L. bicolor* (LB), *H. cylindrosporum* (HC) and *S. indica* (SI). A) Phenotype of Col-0 seedlings. The scale indicates 1 cm. B) Primary root length (cm). C) Number of lateral roots. D) Fresh weight of the entire seedling (mg). Analysis of statistical differences using ANOVA and Tukey’s post hoc test, n=50-100, p<0.05.

### Volatile organic compounds (VOCs) emitted by *L. bicolor, H. cylindrosporum*, and *S. indica* affect WOX5 expression in Arabidopsis

Western blot analysis revealed that *WOX5* expression levels under control conditions (absence of osmotic stress) were increased in the presence of VOCs emitted by *H. cylindrosporum* and, predominantly, *S. indica*, compared to seedlings not exposed to fungal VOCs (Figure 2A). This increase was also observed in confocal microscopy images of the root meristem, thus corroborating the western blot results (Figure 2B).

**Figure 2.**
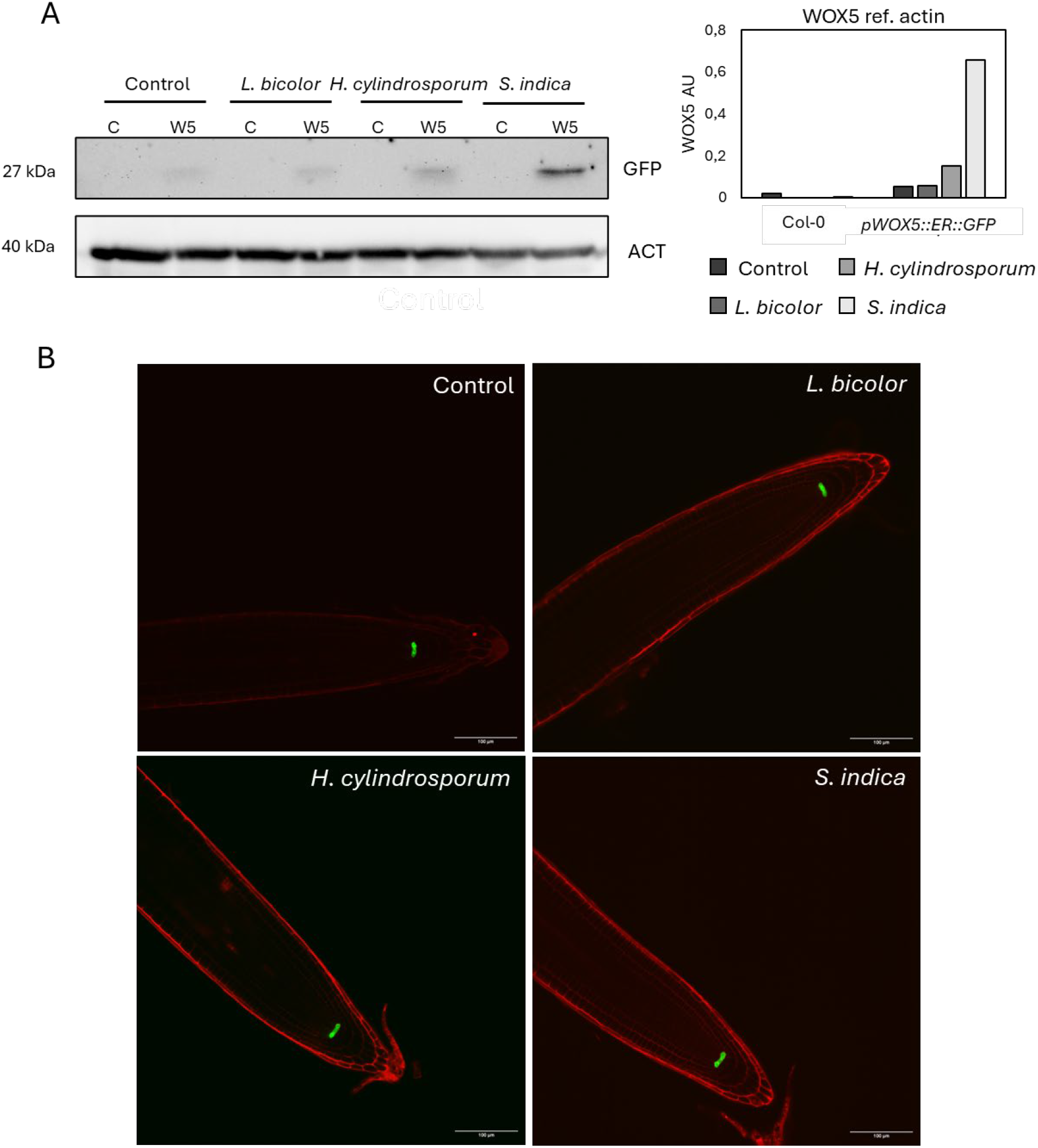
Analysis of root meristem using 6-day-old seedlings, using the *pWOX5::ER:GFP* reporter line in the absence and presence of VOCs from *L. bicolor, H. cylindrosporum*, and *S. indica* under control conditions. A) Analysis of WOX5 by western blot under control conditions (n=3). C is Col-0 and W5 is *pWOX5::ER::GFP*. Normalized with actin accumulation. B) Root meristem observed under a Dragonfly 200 High Speed Confocal microscope. The scale shows 100µm.

Under osmotic stress conditions (100 mM sorbitol), *WOX5* expression levels were elevated in seedlings exposed to VOCs from *L. bicolor* and *H. cylindrosporum* compared to those not exposed to fungal VOCs (Figure 3A). This finding was again confirmed by imaging of the root meristem (Figure 3B).

**Figure 3.**
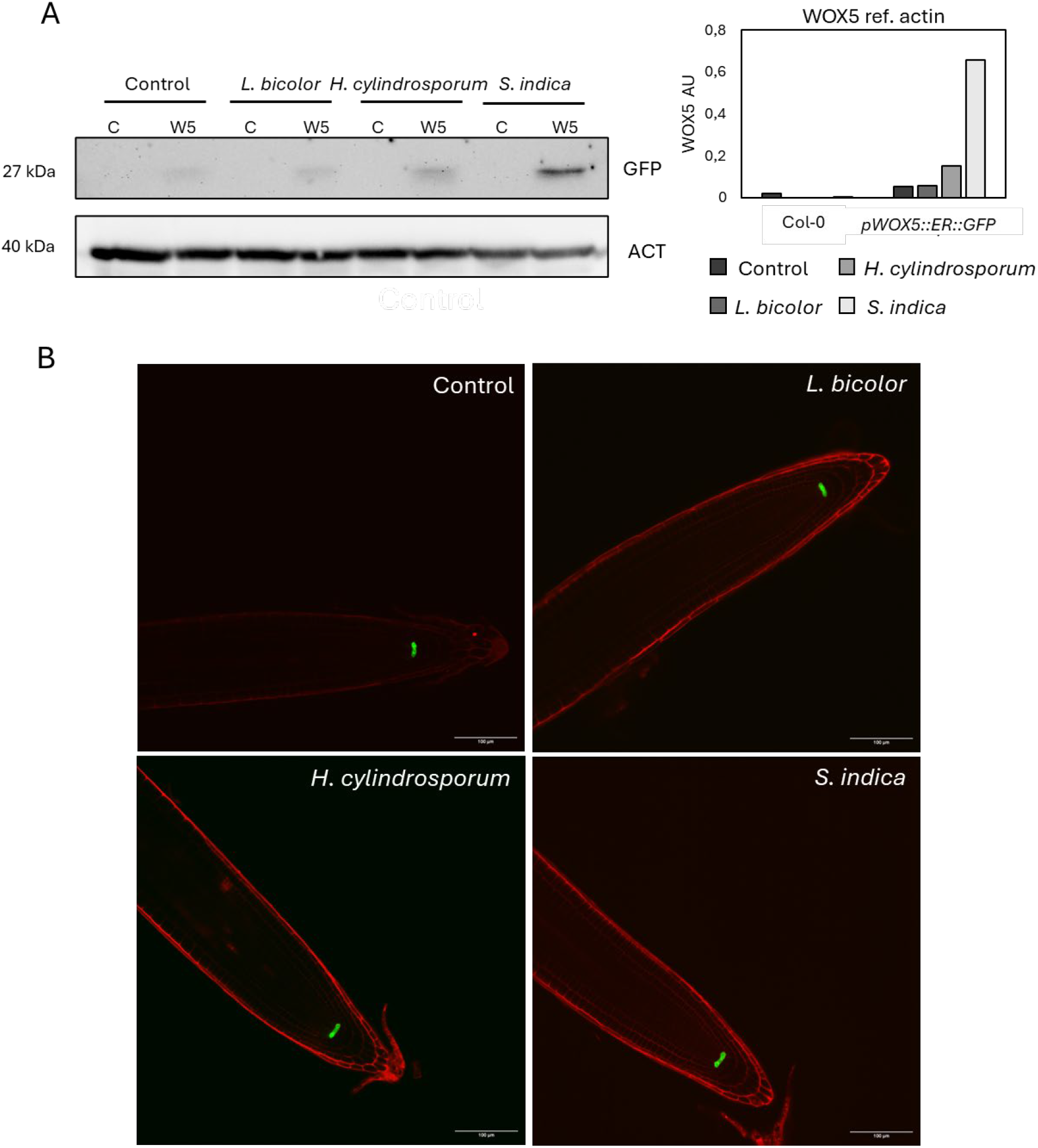
Analysis of root meristem using 6-day-old seedlings, using the *pWOX5::ER:GFP* reporter line in the absence and presence of VOCs from *L. bicolor, H. cylindrosporum*, and *S. indica* under osmotic stress conditions. A) Analysis of WOX5 by western blot under control conditions (n=3). C is Col-0 and W5 is *pWOX5::ER::GFP*. Normalized with actin accumulation. B) Root meristem observed under a Dragonfly 200 High Speed Confocal microscope. The scale shows 100µm. C) Analysis of cell damage in the meristem by quantifying the fluorescence intensity of propidium iodide (PI). Analysis of statistical differences using ANOVA and Tukey’s post hoc test, n=20-30, p<0.05.

Propidium iodide stains cell walls but only penetrates cells that are damaged or dead (Chen & Fluhr, 2018). This technique was employed to assess cellular damage in the root meristem. Seedlings subjected to osmotic stress exhibited greater damage compared to those under control conditions (Figure 3C). Additionally, seedlings exposed to osmotic stress showed less damage both in the absence of fungal VOCs and when exposed to VOCs emitted by *S. indica* (Figure 3C). In contrast, seedlings under osmotic stress and exposed to VOCs from *L. bicolor* and *H. cylindrosporum* appeared the most damaged compared to the control without fungal VOC exposure (Figure 3C).

### Fungal VOCs affects the development of *P. tremuloides* seedlings

To determine the effect of fungal VOCs on other agronomically relevant species, their impact was assessed on the forest species *P. tremuloides*, a typical host of *L. bicolor* (Martin et al., 2016). Under control conditions, seedlings exposed to fungal VOCs exhibited a more developed aerial part and a more complex root system compared to seedlings not exposed to VOCs (Figure 4). Also, the primary root length increased in seedlings exposed to fungal VOCs emitted by any of the three fungi compared to the absence of VOCs (Figure 4A), consistent with observations in Arabidopsis (Figure 1B). However, the average length of lateral roots showed no significant differences under VOC exposure (Figure 4B). The number of lateral roots was higher in seedlings exposed to VOCs from all three fungal species, as in Col-0 ecotype (Figures 1C, Figure 4C). Finally, total fresh weight also increased in response to fungal VOCs (Figure 4D), with a more pronounced increase observed in the aerial parts (Figure 4E) than in the roots (Figure 4F).

**Figure 4.**
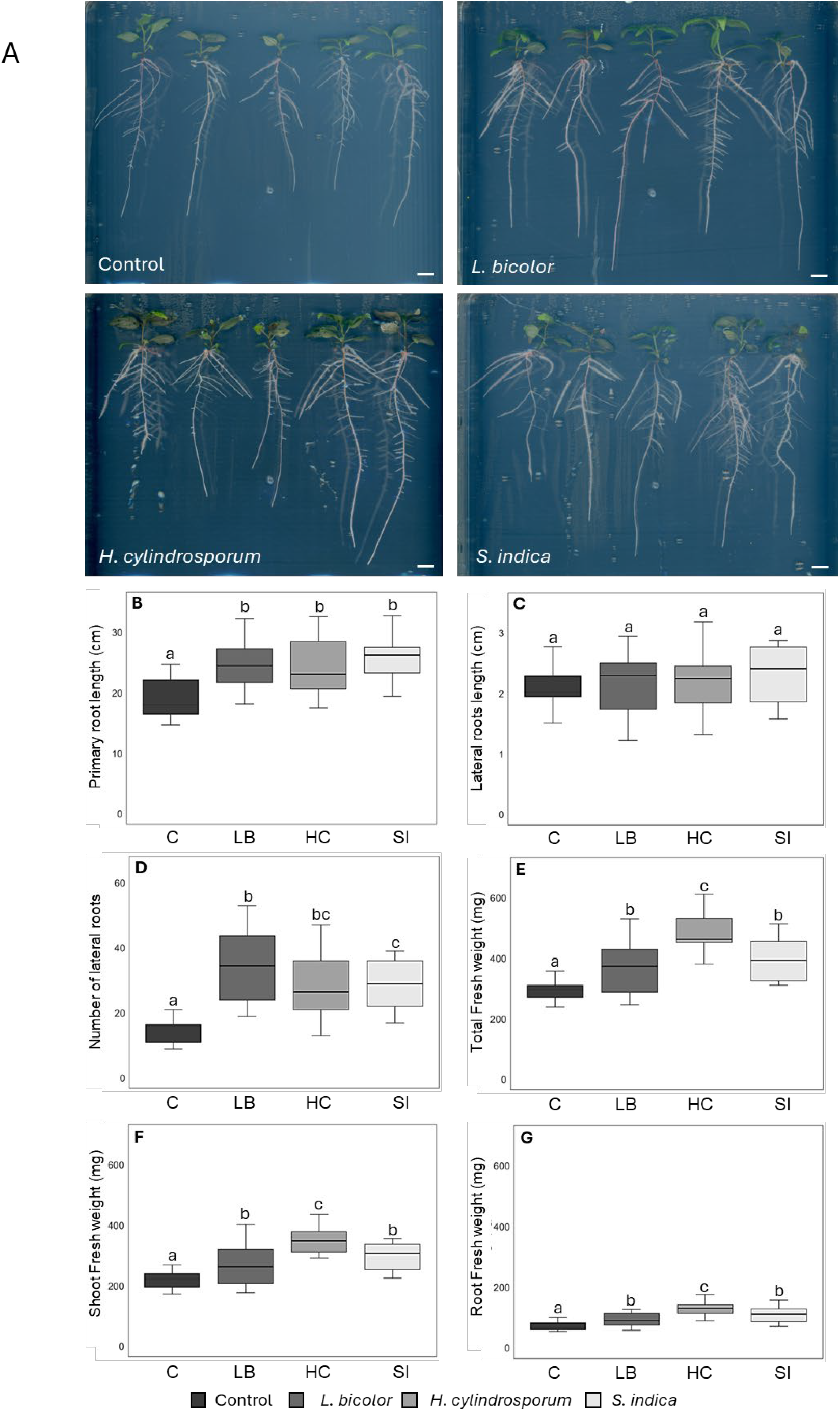
Analysis of the development of 15-day-old seedlings of *P. tremuloides* in the absence and presence of VOCs from *L. bicolor, H. cylindrosporum*, and *S. indica* under controlled conditions. A) Phenotype of *P. tremuloides* seedlings. The scale indicates 1 cm. B) Length of primary root. C) Length of lateral roots. D) Number of lateral roots. E) Total fresh weight. F) Fresh weight of aerial part. G) Fresh weight of root. Analysis of statistical differences using ANOVA test and Tukey’s post hoc test, n=50-100, p<0.05.

### The expression dynamics of *WOX5* and orthologs in *P. tremuloides* are affected by fungal VOCs under control conditions

In *P. tremuloides*, there are two orthologous genes of Arabidopsis *WOX5* (*AT3G11260*), *Potrs008975g13564* (*WOX5*.*1*) and *Potrs019544g21961* (*WOX5*.*2*), identified using the database PlantGenIE (2021). *WOX5*.*1* shows significantly higher expression in roots of *P. tremuloides* seedlings exposed to VOCs emitted by *S. indica* under control conditions compared to seedlings not exposed to fungal VOCs (Figure 5A). However, analysis of *WOX5*.*2* did not reveal significant differences due to high variability among biological replicates, preventing conclusive results (Figure 5B).

**Figure 5.**
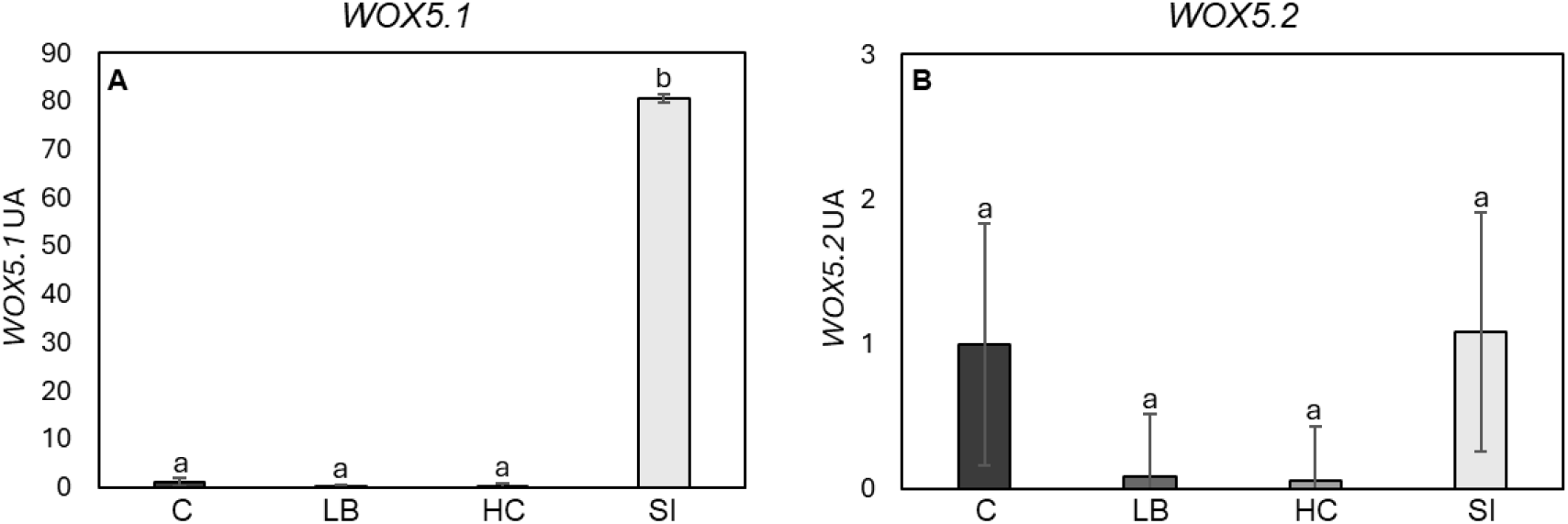
**Analysis of the expression of orthologs of *WOX5* in *P. tremuloides* using RT-qPCR in plants exposed for 15 days to VOCs emitted by *L. bicolor, H. cylindrosporum* and *S. indica* under control conditions, normalized to the control not exposed to fungal VOCs**. A) *Potrs008975g13564* (*WOX5*.*1*). B) Potrs019544g21961 (*WOX5*.*2*).

## Discussion

Ectomycorrhizal fungi such as *L. bicolor* and *H. cylindrosporum* lack the ability to colonize plants from the Brassicaceae family, including Arabidopsis (Smith & Read, 2010), unlike *S. indica* (González Ortega ‐Villaizán et al., 2024; Opitz et al., 2021). Nevertheless, the results obtained in this study revealed that both seedlings from the Col-0 ecotype of *Arabidopsis* and *P. tremuloides* exhibited a significant increase in primary root length as well as in the number of lateral roots when exposed to fungal VOCs released by any of the three fungi tested (Figure 1, 5). Even though Arabidopsis and *P. tremuloides* are highly divergent species with distinct preferential symbiotic fungi interactions (González Ortega ‐Villaizán et al., 2024; Opitz et al., 2021; Martin et al., 2016; Smith & Read, 2010), root modification induced by fungal VOCs occurred in both species to a comparable extent.

Previous studies had already reported that VOCs emitted by *L. bicolor* promote lateral root development in both *Arabidopsis* and *Populus tremula × alba* (Ditengou et al., 2015; Felten et al., 2010). Similar effects of *S. indica* VOCs on root system architecture have also been described in Arabidopsis and in additional plant species such as *Ocimum basilicum* (Fraj & Werbrouck, 2023; Venneman et al., 2020). The results obtained in the present work showed that exposure of Arabidopsis and *P. tremuloides* roots to the VOCs released by *L. bicolor, H. cylindrosporum*, or *S. indica* resulted in the same pattern of growth responses. These findings therefore suggest that the main molecular signaling pathways mediating root modification might be conserved, regardless of the plant species exposed to fungal VOCs or the fungal identity of the VOC-producing organism, at least within ectomycorrhizal and endophytic symbionts. Consequently, the research was continued with the aim of investigating these molecular signaling mechanisms in greater detail.

The expression of the transcription factor *WOX5* allows stem cells surrounding the niche to remain in an undifferentiated state while maintaining their proliferative capacity, thereby sustaining root growth (Burkart et al., 2022; Pardal & Heidstra, 2021; Sarkar et al., 2007). Consequently, *WOX5* induction leads to an altered meristem, characteristically broadened, with cells exhibiting reduced differentiation capacity, ultimately affecting both root growth and curvature (Burkart et al., 2022; Savina et al., 2020).

The effects of fungal VOCs on *WOX5* expression in Arabidopsis differ under control and osmotic stress conditions (Figures 2, 3). Under control conditions, *WOX5* expression was primarily upregulated in the presence of *S. indica* (Figure 2). In contrast, under osmotic stress, *WOX5* expression increased in response to VOCs released by *L. bicolor* and *H. cylindrosporum* (Figure 3). These findings indicate that VOCs-mediated signaling in the root meristem differs between ectomycorrhizal and endophytic fungi, and that it directly depends on the presence or absence of osmotic stress. The effects on *WOX5* are expected not only to alter meristematic activity but also to influence overall root system architecture, promoting primary root growth while suppressing lateral root initiation because of disrupted auxin flux (Savina et al., 2020). However, the phenotypic effects observed in this study revealed neither primary root inhibition nor lateral root suppression, even in conditions where *WOX5* expression was enhanced.

To visualize cellular damage under osmotic stress, propidium iodide staining was employed, as this compound is unable to penetrate intact membranes and thus selectively marks damaged or dead cells. Given that cells exposed to osmotic stress undergo dehydration, accumulate reactive oxygen species (ROS), and experience changes in membrane structure, propidium iodide was able to infiltrate these cells more extensively compared to non-stressed controls (Chen & Fluhr, 2018). In Arabidopsis under osmotic stress conditions (Figure 3C), stronger staining revealed that root meristems exposed to VOCs emitted by *L. bicolor* and *H. cylindrosporum* displayed greater cellular permeability. This observation coincided with *WOX5* upregulation, suggesting altered meristems with reduced differentiation capacity (Savina et al., 2020).

The ortholog of *WOX5* in *Populus tremuloides, Potrs008975g13564* (*WOX5*.*1*), has previously been described as the active form in *Populus trichocarpa* and *Populus tomentosa*. In these species, induction of the *WOX5* ortholog promotes lateral root organogenesis while simultaneously inhibiting their growth by repressing the expression of differentiation-associated genes, through auxin transport and signaling pathways. Moreover, *WOX5* has been implicated in adventitious root formation as well as in root regeneration in poplar (Li et al., 2018). The results obtained in this work suggest that under control conditions the expression dynamics of *Potrs008975g13564 (WOX5*.*1)* resemble those observed in Arabidopsis, with increased expression in the presence of VOCs emitted by *S. indica* (Figure 5A). This further supports the notion that the effects of fungal VOCs on *WOX5* regulation are conserved across plant species. The evolutionary conservation of these expression dynamics highlights the central role of *WOX5* in mediating plant responses to fungal VOCs.

## Material and methods

### Plant material and fungal strains

*Arabidopsis thaliana* (L.) Heynh ecotypes Columbia (Col-0) was the genetic background used in this work. Additionally, the reporter line *pWOX5::ER::GFP* was employed (Blilou et al., 2005). *Populus tremuloides*, commonly known as aspen, is the most widespread tree in North America (Landhäusser et al., 2019; Perala et al., 1990). It is characterized by being one of the few species of poplar that is commercially available and reproduces by seed for exploitation. Its economic importance is very significant, both in the timber industry and in paper production, as well as for its ecological role. This poplar is a pioneer species in the colonization of soils that have suffered fires or other types of disturbances, and through vegetative reproduction from the root, it can occupy large areas of land. In addition, it prevents soil erosion, and, due to its great ecological plasticity, it can adapt to different environments (Perala et al., 1990).

*Laccaria bicolor* (Maire) P. D. Orton is an ectomycorrhiza of the Tricholomataceae family. The subculture S238N was obtained from the Institute National de la Recherche Agronomique (INRAE) (Bertaux et al., 2003). *Hebeloma cylindrosporum* (Romagnesi) is an ectomycorrhizal fungus of the Hymenogastraceae family, obtained from the INRA in Monpellier, and the endophytic fungi *Serendipita indica* Verma, Varma, Kost, Rexer & Franken (Varma et al., 1999; Verma et al., 1998) was obtained from the University of Murcia. The different fungal strains were cultivated in Pachelewski solid medium (Pachlewski & Pachlewska, 1974) in petri dishes plates.

### Growth conditions and plant fungus co-culture

Arabidopsis seeds were surface sterilized in 75 % (v/v) sodium hypochlorite and 0.01 % (v/v) Triton X-100 for 5 min and washed three times in sterile water before sowing. Seeds were stratified for 24h at 4 °C. With respect to *P. tremuloides*, seeds were superficially cleaned using a solution of 10% (v/v) sodium hypochlorite and 0.01 % (v/v) Triton X-100 for 10 minutes with agitation, after which they were washed three times with sterile distilled water. The seeds were stratified 24 hours at 4 °C.

*Arabidopsis thaliana* seedlings were grown in 120 x 120 mm square plates (Fisher Scoientific) filled with MS medium (Murashige, 1962), into which three round plates measuring 35mm (FALCON, A Corning Brand) and filled with Pachlewski P20 medium (Pachlewski & Pachlewska, 1974) were added. For the osmotic stress condition, the plates were prepared in the same way as explained above, but with MS medium supplemented with sorbitol (100 mM). 40 seeds of the Col-0 ecotype and *pWOX5::ER::GFP* were sawn and the plates were sealed with a double layer of Parafilm and left in a Fitoclima 600 Aralab climate chamber, under controlled conditions of 50-60% humidity, 16:8 hours (day:night), 22 °C and150 μE m^−2^ s^−1^. The plates were protected from light with a black cover so that only the Arabidopsis area was exposed to light, and the root and fungi remained in darkness. When the seedlings were 10 days old, they were scanned using an EPSON V850 Pro system and their fresh weight root length and number of secondary roots were determined.

For the co-cultivation assays with fungi and *P. tremuloides*, a similar system was used but with larger plates (245×245mm, Corning). Within them, four round plates measuring 55mm (Corning) were inserted. Pachlewski P20 medium (Pachlewski & Pachlewska, 1974) was added to the round plates, and McCown or Woody Plant Medium (Lloyd & McCown, 1980) to the square one. Nine-day-old *P. tremuloides* seedlings (germinated in McCown medium) were transferred to the square plate six days after sowing the fungus on the small plates with Pachlewski P20. The plates were sealed with a double layer of Parafilm and incubated in a Fitoclima 600 Aralab climate chamber, under conditions of 50-60% humidity, with a photoperiod of 16 hours of light at 22 °C and 8 hours of darkness at 18°C, at a light intensity of 300 μE m^−2^ s^−1^. 15 days after seed sowing fresh weights were determined after the plates were scanned using an EPSON V850 Pro system and the images analyzed with the SmartRoot software to determine primary and lateral root length and the number of lateral roots.

### Western blotting

Total protein for western blot analysis was extracted from the root tips of 6-day-old Arabidopsis seedlings. The tissue was ground in a mortar with liquid nitrogen and subsequently homogenized in extraction buffer (150 mM NaCl, 0.25% NP-40, 100 mM Tris-HCl, 1 mM phenylmethylsulphonyl fluoride, and 1X EDTA-free Complete Protease Inhibitor Cocktail (Sigma)) using a Silamat S6 for 10 seconds, followed by centrifugation at 13,000 rpm for 15 minutes at 4 °C. Protein concentration was measured using the Bradford method with the Bio-Rad Protein Assay kit (Bradford, 1976).

Fifty micrograms of total protein were loaded per well for SDS–polyacrylamide gel electrophoresis using Tris–glycine–SDS running buffer. Proteins were transferred electrophoretically onto Immobilon-P polyvinylidene difluoride (PVDF) membranes (Millipore) with the Trans-Blot Turbo system (Bio-Rad). Membranes were blocked in Tris-buffered saline containing 0.1% Tween 20 and 5% Blocking Agent, then incubated with primary antibodies diluted in blocking buffer.

For detection, Living Colors® A.v. Monoclonal Antibody (JL-8, CLONTECH) was used for anti-GFP, followed by either anti-rabbit IgG or anti-mouse IgG secondary antibodies (Fisher Scientific NA934 and NA931, respectively; both 1:10,000 dilution). Chemiluminescent signals were developed with SuperSignal West Femto Maximum Sensitivity Substrate (Thermo Scientific) and imaged using the ChemiDoc MP Imaging System (Bio-Rad). Band intensities were quantified with ImageJ software.

### Quantitative reverse transcriptase–PCR analysis

Total RNA was extracted from roots of 10 day-old seedlings. Briefly, frozen tissue was powdered with mortar and as conserved at -80ºC until use. RNA was extracted with RNeasy Plant Mini Kit (QIAGEN) and was precipitated in RNAse free water. RNA concentration was measured in NanoDrop 1000 (Thermo Scientific). Retrotrascriptation reaction was performed by iScript Select cDNA Synthesis kit (BioRad). Later, a RT-PCR were performed using a StepOnePlus Real-Time PCR system (Thermo Fisher Scientific). Amplification was carried out with SYBR FAST Rox High (KAPA system) according to the manufacturer’s instructions. The thermal profile for SYBR Green real-time PCR was 50 °C for 2 min, 95 °C for 10 min, followed by 40 cycles of 95 °C for 15 s and 60 °C for 1 min.

To generate the standard curves, cDNA isolated from *Arabidopsis* seedlings was serially diluted 10 × and aliquots of the dilutions were used in standard real-time PCRs. Each value determination was repeated three times, to ensure the slope of the standard curves and to determine the s.d. The concentration of unknown samples was calculated with the ABI-Prism 7000 SDS software, which created threshold cycle values (*C*_t_) and extrapolated relative levels of PCR product from the standard curve.

## References

Bertaux, J., Schmid, M., Prevost-Boure, N. C., Churin, J. L., Hartmann, A., Garbaye, J., & Frey-Klett, P. (2003). In Situ Identification of Intracellular Bacteria Related to Paenibacillus spp. in the Mycelium of the Ectomycorrhizal Fungus Laccaria bicolor S238N. Applied and Environmental Microbiology, 69(7), 4243–4248.

Blilou, I., Xu, J., Wildwater, M., Willemsen, V., Paponov, I., Friml, J., Heidstra, R., Aida, M., Palme, K., & Scheres, B. (2005). The PIN auxin efflux facilitator network controls growth and patterning in Arabidopsis roots. Nature, 433(7021), 39–44.

Burkart, R. C., Strotmann, V. I., Kirschner, G. K., Akinci, A., Czempik, L., Dolata, A., Maizel, A., Weidtkamp-Peters, S., & Stahl, Y. (2022). PLETHORA-WOX5 interaction and subnuclear localization control Arabidopsis root stem cell maintenance. EMBO Reports, 23(6).

Chen, T., & Fluhr, R. (2018). Singlet Oxygen Plays an Essential Role in the Root’s Response to Osmotic Stress. Plant Physiology, 177(4), 1717–1727.

Cho, H. J., Park, K. H., Park, M. S., Cho, Y., Kim, J. S., Seo, C. W., Oh, S.-Y., & Lim, Y. W. (2021). Determination of Diversity, Distribution and Host Specificity of Korean Laccaria Using Four Approaches. Mycobiology, 49(5), 461–468.

Cope, K. R., Bascaules, A., Irving, T. B., Venkateshwaran, M., Maeda, J., Garcia, K., Rush, T. A., Ma, C., Labbé, J., Jawdy, S., Steigerwald, E., Setzke, J., Fung, E., Schnell, K. G., Wang, Y., Schleif, N., Bücking, H., Strauss, S. H., Maillet, F., … Ané, J.-M. (2019). The Ectomycorrhizal Fungus Laccaria bicolor Produces Lipochitooligosaccharides and Uses the Common Symbiosis Pathway to Colonize Populus Roots. The Plant Cell, 31(10), 2386–2410.

Ditengou, F. A., Müller, A., Rosenkranz, M., Felten, J., Lasok, H., van Doorn, M. M., Legué, V., Palme, K., Schnitzler, J.-P., & Polle, A. (2015). Volatile signalling by sesquiterpenes from ectomycorrhizal fungi reprogrammes root architecture. Nature Communications, 6(1), 6279.

Drijber, R. A., & McPherson, M. R. (2021). Mycorrhizal symbioses. In Principles and Applications of Soil Microbiology (pp. 303–325). Elsevier.

El Jaddaoui, I., Rangel, D. E. N., & Bennett, J. W. (2023). Fungal volatiles have physiological properties. Fungal Biology, 127(7–8), 1231–1240.

Felten, J., Legué, V., & Anicet Ditengou, F. (2010). Lateral root stimulation in the early interaction between Arabidopsis thaliana and the ectomycorrhizal fungus Laccaria bicolor. Plant Signaling & Behavior, 5(7), 864–867.

Fraj, H., & Werbrouck, S. P. O. (2023). Constant and Intermittent Contact with the Volatile Organic Compounds of Serendipita indica Alleviate Salt Stress In Vitro Ocimum basilicum L. Applied Sciences, 13(3), 1776.

Gautam, A. K., & Avasthi, S. (2019). Fungal endophytes: potential biocontrol agents in agriculture. In Role of Plant Growth Promoting Microorganisms in Sustainable Agriculture and Nanotechnology (pp. 241–283). Elsevier.

González Ortega-Villaizán, A., King, E., Patel, M. K., Pérez-Alonso, M., Scholz, S. S., Sakakibara, H., Kiba, T., Kojima, M., Takebayashi, Y., Ramos, P., Morales-Quintana, L., Breitenbach, S., Smolko, A., Salopek-Sondi, B., Bauer, N., Ludwig-Müller, J., Krapp, A., Oelmüller, R., Vicente-Carbajosa, J., & Pollmann, S. (2024). The endophytic fungus Serendipita indica affects auxin distribution in Arabidopsis thaliana roots through alteration of auxin transport and conjugation to promote plant growth. Plant, Cell & Environment, 47(10), 3899–3919.

Inamdar, A. A., Morath, S., & Bennett, J. W. (2020). Fungal Volatile Organic Compounds: More Than Just a Funky Smell? Annual Review of Microbiology, 74(1), 101–116.

Jalali, F., Zafari, D., & Salari, H. (2017). Volatile organic compounds of some Trichoderma spp. increase growth and induce salt tolerance in Arabidopsis thaliana. Fungal Ecology, 29, 67–75.

Kong, X., Liu, G., Liu, J., & Ding, Z. (2018). The Root Transition Zone: A Hot Spot for Signal Crosstalk. Trends in Plant Science, 23(5), 403–409.

Kumar, P., Devi, P., & Dey, S. R. (2021). Fungal volatile compounds: a source of novel in plant protection agents. In Volatiles and Metabolites of Microbes (pp. 83–104). Elsevier.

Landhäusser, S. M., Pinno, B. D., & Mock, K. E. (2019). Tamm Review: Seedling-based ecology, management, and restoration in aspen (Populus tremuloides). Forest Ecology and Management, 432, 231–245.

Li, J., Zhang, J., Jia, H., Liu, B., Sun, P., Hu, J., Wang, L., & Lu, M. (2018). The WUSCHEL-related homeobox 5a (PtoWOX5a) is involved in adventitious root development in poplar. Tree Physiology, 38(1), 139–153.

Li, N., & Kang, S. (2018). Do volatile compounds produced by Fusarium oxysporum and Verticillium dahliae affect stress tolerance in plants? Mycology, 9(3), 166–175.

Marmeisse, R., Guidot, A., Gay, G., Lambilliotte, R., Sentenac, H., Combier, J.-P., Melayah, D., Fraissinet-Tachet, L., & Debaud, J. C. (2004). Hebeloma cylindrosporum– a model species to study ectomycorrhizal symbiosis from gene to ecosystem. New Phytologist, 163(3), 481–498.

Martin, F., Aerts, A., Ahrén, D., Brun, A., Danchin, E. G. J., Duchaussoy, F., Gibon, J., Kohler, A., Lindquist, E., Pereda, V., Salamov, A., Shapiro, H. J., Wuyts, J., Blaudez, D., Buée, M., Brokstein, P., Canbäck, B., Cohen, D., Courty, P. E., … Grigoriev, I. V. (2008). The genome of Laccaria bicolor provides insights into mycorrhizal symbiosis. Nature, 452(7183), 88–92.

Martin, F., Kohler, A., Murat, C., Veneault-Fourrey, C., & Hibbett, D. S. (2016). Unearthing the roots of ectomycorrhizal symbioses. Nature Reviews Microbiology, 14(12), 760–773.

Opitz, M. W., Daneshkhah, R., Lorenz, C., Ludwig, R., Steinkellner, S., & Wieczorek, K. (2021). Serendipita indica changes host sugar and defense status in Arabidopsis thaliana: cooperation or exploitation? Planta, 253(3), 74.

Pardal, R., & Heidstra, R. (2021). Root stem cell niche networks: it’s complexed! Insights from Arabidopsis. Journal of Experimental Botany, 72(19), 6727–6738.

Perala, D. A., Burns, R. M., & Honkala, B. (1990). Populus tremuloides Michx.-Quaking Aspen. Silvics of North America: Hardwoods; Burns, RM, Honkala, BH, Eds, 555–569.

Ruytinx, J., Miyauchi, S., Hartmann-Wittulsky, S., de Freitas Pereira, M., Guinet, F., Churin, J.-L., Put, C., Le Tacon, F., Veneault-Fourrey, C., Martin, F., & Kohler, (2021). A Transcriptomic Atlas of the Ectomycorrhizal Fungus Laccaria bicolor. Microorganisms, 9(12), 2612.

Saleem, S., Sekara, A., & Pokluda, R. (2022). Serendipita indica—A Review from Agricultural Point of View. In Plants (Vol. 11, Issue 24). MDPI.

Sarkar, A. K., Luijten, M., Miyashima, S., Lenhard, M., Hashimoto, T., Nakajima, K., Scheres, B., Heidstra, R., & Laux, T. (2007). Conserved factors regulate signalling in Arabidopsis thaliana shoot and root stem cell organizers. Nature, 446(7137), 811–814.

Savina, M. S., Pasternak, T., Omelyanchuk, N. A., Novikova, D. D., Palme, K., Mironova, V. V., & Lavrekha, V. V. (2020). Cell Dynamics in WOX5-Overexpressing Root Tips: The Impact of Local Auxin Biosynthesis. Frontiers in Plant Science, 11.

Singh, J., Singh, P., Vaishnav, A., Ray, S., Rajput, R. S., Singh, S. M., & Singh, H. A.(2021). Belowground fungal volatiles perception in okra (Abelmoschus esculentus) facilitates plant growth under biotic stress. Microbiological Research, 246, 126721.

Smith, S. E., & Read, D. J. (2010). Mycorrhizal symbiosis. Academic press.

Stahl, Y., & Simon, R. (2005). Plant stem cell niches. The International Journal of Developmental Biology, 49(5–6), 479–489.

Su, Y.-H., Liu, Y.-B., & Zhang, X.-S. (2011). Auxin-Cytokinin Interaction Regulates Meristem Development. Molecular Plant, 4(4), 616–625.

Varma, A., Savita Verma, Sudha, Sahay, N., Bütehorn, B., & Franken, P. (1999). Piriformospora indica, a Cultivable Plant-Growth-Promoting Root Endophyte. Applied and Environmental Microbiology, 65(6), 2741–2744.

Venneman, J., Vandermeersch, L., Walgraeve, C., Audenaert, K., Ameye, M., Verwaeren, J., Steppe, K., Van Langenhove, H., Haesaert, G., & Vereecke, D. (2020). Respiratory CO2 Combined With a Blend of Volatiles Emitted by Endophytic Serendipita Strains Strongly Stimulate Growth of Arabidopsis Implicating Auxin and Cytokinin Signaling. Frontiers in Plant Science, 11, 544435.

Verma, S., Varma, A., Rexer, K.-H., Hassel, A., Kost, G., Sarbhoy, A., Bisen, P., Bütehorn, B., & Franken, P. (1998). Piriformospora indica, gen. et sp. nov., a new root-colonizing fungus. Mycologia, 90(5), 896–903.

